# The within-host viral kinetics of SARS-CoV-2

**DOI:** 10.1101/2020.02.29.965418

**Authors:** Chentong Li, Jinhu Xu, Jiawei Liu, Yicang Zhou

## Abstract

In this work, we use a within-host viral dynamic model to describe the SARS-CoV-2 kinetics in host. Chest radiograph score data are used to estimate the parameters of that model. Our result shows that the basic reproductive number of SARS-CoV-2 in host growth is around 3.79. Using the same method we also estimate the basic reproductive number of MERS virus is 8.16 which is higher than SARS-CoV-2. The PRCC method is used to analyze the sensitivities of model parameters and the drug effects on virus growth are also implemented to analyze the model.

## 1 Introduction

At the end of 2019, a new type of coronavirus, SARS-CoV-2, began to threaten the people in China, especially in Hubei province. As of March.3, 2020, 80302 individuals have been confirmed to be infected by this virus in China, including 2946 deaths. To mitigate the spread of the virus, the Chinese Government has progressively implemented metropolitan-wide quarantine in Wuhan and several nearby cities from Jan 23–24, 2020 [18]. The virus has also spread to several other countries, such as the Republic of Korea, Japan, Italy, and USA. Around the world, many new deaths are reported every day.

Chest CT is used to assess the severity of lung involvement in COVID-19 [14], which is the name of that new coronavirus caused disease [17]. Before Feb.13, the new cases in China were confirmed by nucleic acid testing. After that day, the diagnostic criterion has been improved, composed of not only the nucleic acid testing but also the CT test. The destructed pulmonary parenchyma and the resulting inflammation can be reflected from the chest radiograph [13]. Chest radiograph score method is a useful way to quantify the destruction of pulmonary parenchyma and in this study, our data set is based on this.

Several works are [7,18] nowcasting and forecasting the number of confirmed cases of COVID-19. Meanwhile, some works [6,15] overview the characteristics, exposure history, and illness timelines of confirmed cases. All of these works focus on population dynamics of COVID-19, while, the works about the viral dynamics in host are rare. In this work, we use the within-host viral dynamic model [1,11] to describe the SARS-CoV-2 kinetics in host and the parameters of that model are estimated via the chest radiograph score. Moreover, using the CT score data of MERS that shown in Oh et. al [13], the paramters of MERS virus are also estimated as a comparision group. All of the results are shown in the third section.

## 2 Method

We use the following ordinary differential equation model to simulate the coronavirus within-lung growth:

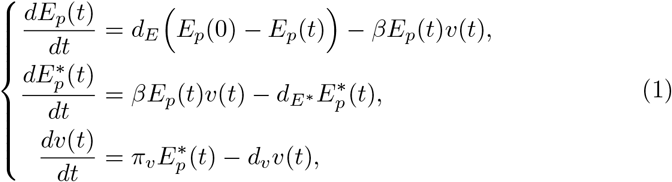

where 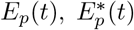 and *v*(*t*) are the number of uninfected pulmonary epithelial cells, infected pulmonary epithelial cells and the virus. *β* is the infection rate of virus, *π_v_* is the virus production rate, *E_p_* (0) is the initial value of uninfected epithelial cells, and the term *d_E_E_p_*(0) assumes a constant regeneration of uninfected epithelial cells. *d_E_*, *d_E_*\* and *d_v_* are the death rate of uninfected pulmonary epithelial cells, infected pulmonary epithelial cells and the virus, respectively. The death rate of *E_p_* (*t*) is the natural clearance rate of pulmonary epithelial cells, while, *d_E_*\* and *d_v_* are the combination of the natural clearance rate and elimination by the immunity system. This model was also used to describe the within-lung infection process of flu virus [5, 11].

By the definition of generation matrix of basic reproduction number *R_0_* [3], the *R_0_* of model (1) can be written as,

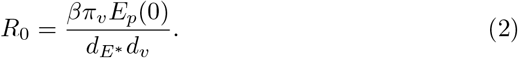

This number is an important value to measure whether the epidemic or species could die out or not [16]. In this study, the sensitivity analysis result of *R*_0_ is shown in the next section.

In this study, the chest radiograph score data from serve patients (with high chest radiograph scores) are collected from the work of Pan et. al [14] and Oh et. al [13] Our estimation is based on these two data sets. We consider the chest radiograph score as a way to reflect the infected pulmonary epithelial cells [13,14], which is also the target attacked by the immune cell [12]. Thus by the Poisson distribution, the likelihood that used to extimate parameters can be written as,

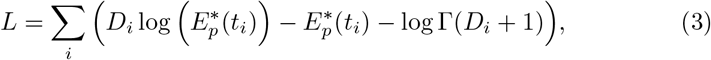

where *D_i_* is the smoothed chest radiograph score of the serve patient at time *t_i_*, 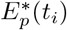 is the solution of 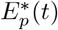 at time *t_i_*, and logΓ(*x*) is log-gamma function, respectively. The prior distributions of the parameters of model (1) are based on the previous works [5,10].

## 3 Results

Figure.1 shows the fitted result of our model and Table.1 summarizes the estimation results of the parameters and *R*_0_ of our model (1). We estimate that the death rate of these two virus are 5.36 (COVID-19) and 4.64 (MERS) per day, which is larger than the clearance rate of the virus on the outside surface [4]. This result shows the immune system can clear the virus directly. The estimation result also shows that the *R*_0_ of SARS-CoV-2 in serve patients is around the mean value 3.79 which is lower than that of MERS virus (8.16). These results illustrate that the immune system can not clear the virus effectively at the beginning time of symptom onset. By these estimated parameters of COVID-19, the solutions of model (1) are illustrated in Figure .2.

**Figure 1:**
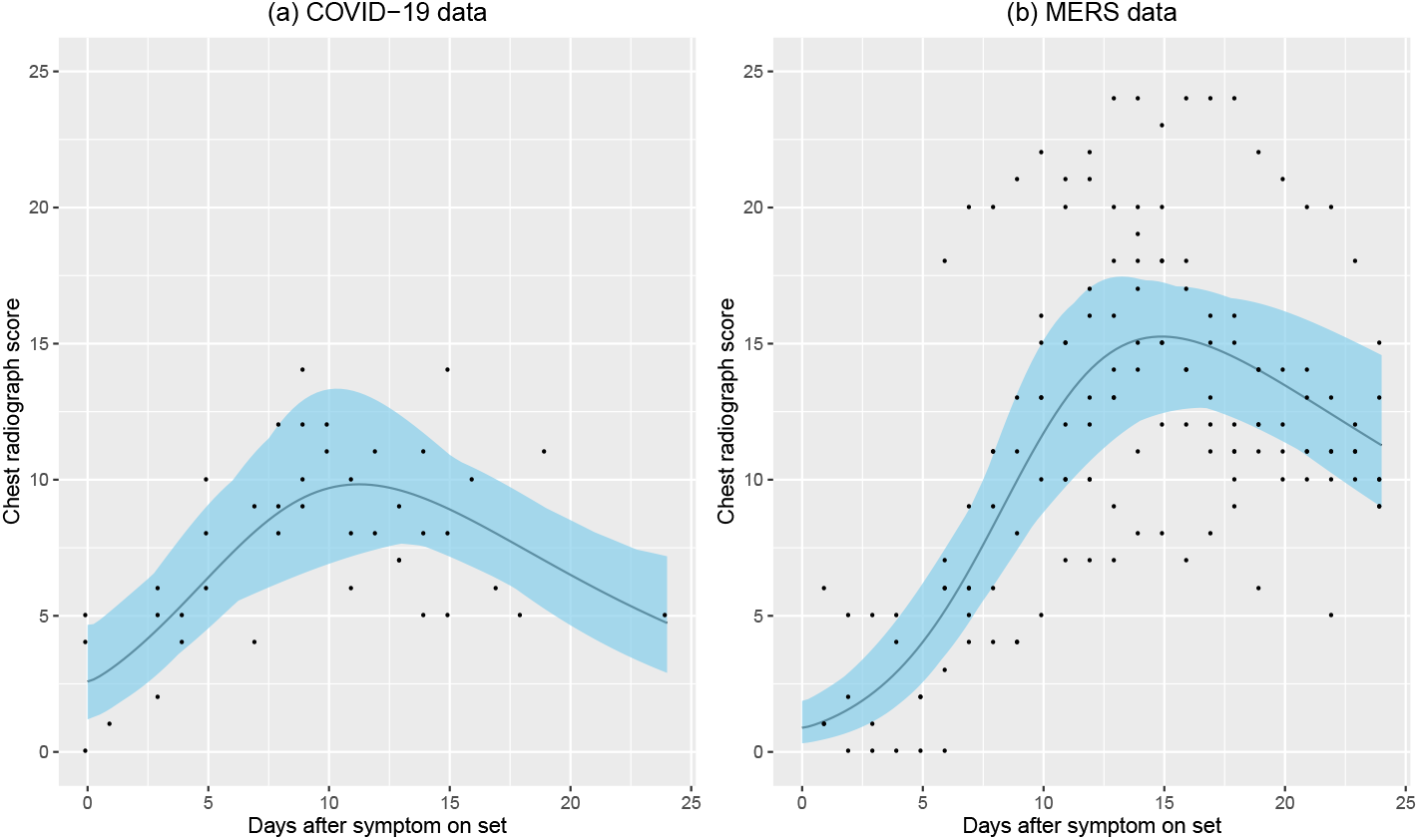
The chest radiograph score data and fitted result of (a) COVID-19, and (b) MERS. The dark line is the mean value of fitted result, while, the 95% confidence interval is shown in blue. The black points are the chest radiograph score data.

**Table 1:** Model parameters

| Parameter | Description (units) | Mean value<br>of COVID-19 (std) | Mean value<br>of MERS<br>(std) | Source |
| --- | --- | --- | --- | --- |
| $E_p(0)$ | The initial value of uninfected epithelial cells (score) | $25 - E_p^*(0)$ | $25 - E_p^*(0)$ | [13, 14] |
| $E_p^*(0)$ | The initial value of infected epithelial cells (score) | 2.59 (0.61) | 0.89 (0.28) | MCMC |
| $v(0)$ | The initial value of virus(score) | 0.061<br>(0.0503) | 0.0075<br>(0.0099) | MCMC |
| $\pi_v$ | Virus production rate per infected epithelial cells ($\text{day}^{-1}$) | 0.24 (0.22) | 0.15 (0.16) | MCMC |
| $\beta$ | Infection rate of epithelial cells by virus ($\text{day}^{-1}\text{score}^{-1}$) | 0.55 (0.55) | 1.28 (1.37) | MCMC |
| $d_E$ | Death rate of epithelial cells ($\text{day}^{-1}$) | $10^{-3}$ | $10^{-3}$ | [5] |
| $d_{E^*}$ | Death rate of infected epithelial cells ($\text{day}^{-1}$) | 0.11 (0.01) | 0.056<br>(0.0053) | MCMC |
| $d_v$ | Death rate of virus ($\text{day}^{-1}$) | 5.36 (6.42) | 4.64 (0.89) | MCMC |
| $R_0$ | Basic reproduction number | 3.79 (0.54) | 8.16 (1.13) | MCMC |

**Figure 2:**
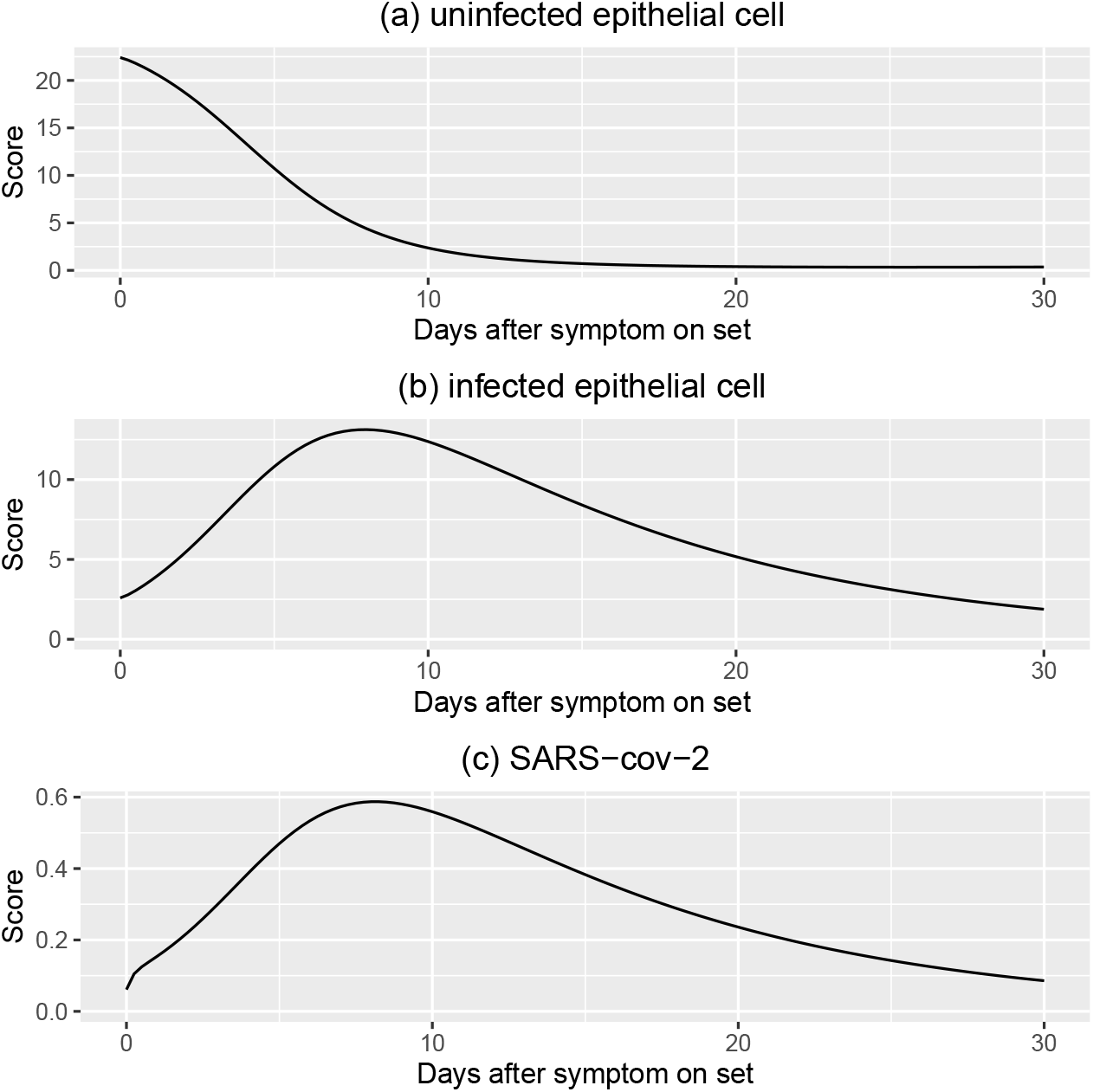
The solutions of model (1) with the parameters of COVID-19 that shown in Table.1.

The partial rank correlation coefficient (PRCC) method [9] is used to do the sensitivity analysis of *R*_0_. The result (Figure .3) shows that the parameters *E_p_*(0), *β*, *π_v_*, *d_E_*\*, and *d_v_* have almost the same level of influence on *R*_0_. Parameters *d_E_*\*, and *d_v_* have negative correlations with *R*_0_, while *E_p_*(0), *β*, *π_v_* have positive correlations. The positive correlation of *E_p_*(0) with *R*_0_ may give a explanation on why the babys, who have small amount of pulmonary epithelial cells, are not likely to get infected and have lower mortality ([15] Table 1).

**Figure 3:**
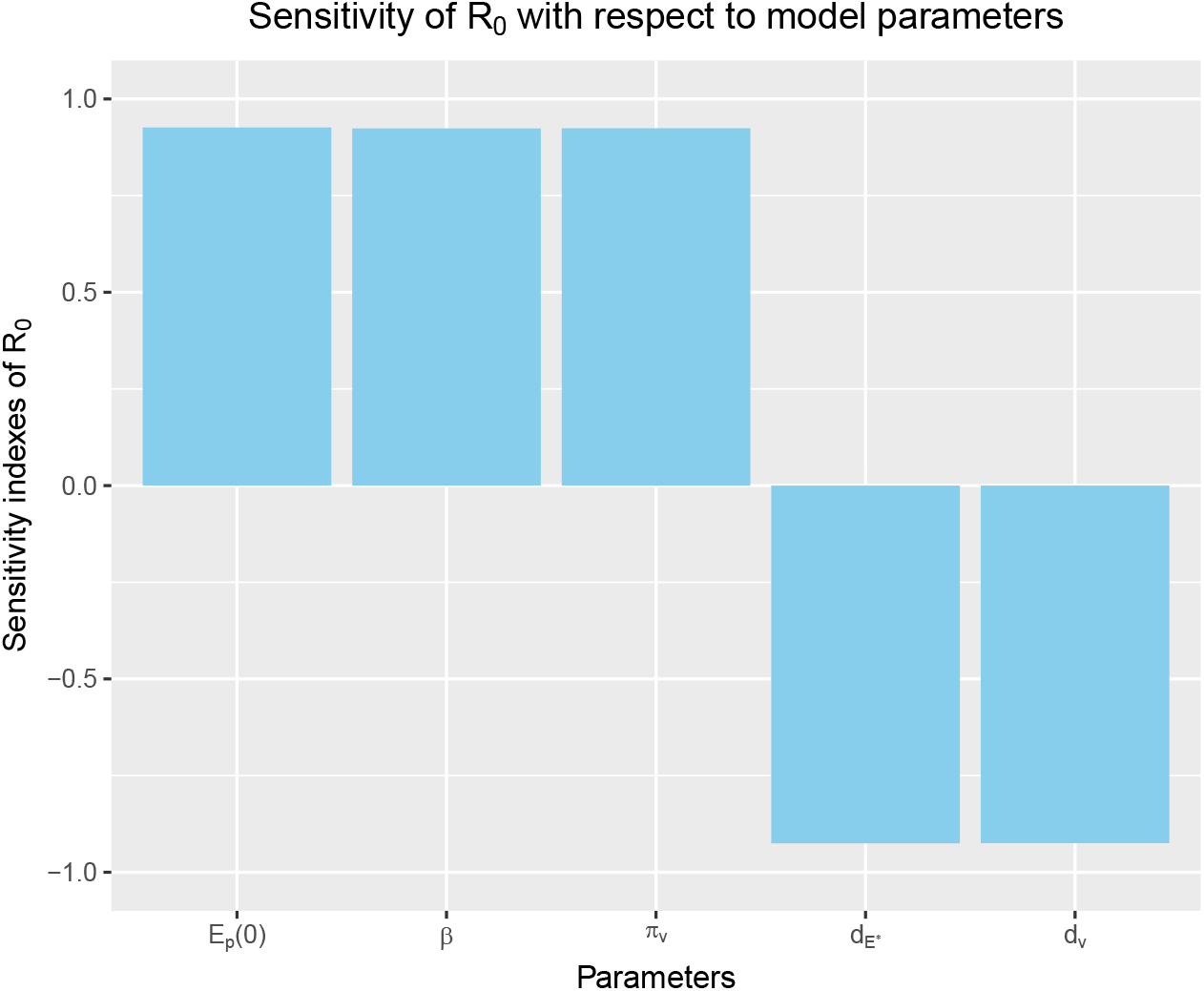
Sensitivity of *R*_0_ with respect to *E_p_*(0), *β*, *π_v_*, *d_E_*\*, and *d_v_*. The samples of parameters are taken from the uniform distribution *U*(0.8*p*, 1.2*p*), where *p* is used to illustrate the mean value of parameters of COVID-19 that shown in Table.1.

Drug therapy is also a topic discussed widely on viral dynamics [1]. For the coronavirus there are some drugs that may have a positive influence on decreasing the virus load [8]. In this study, we assume that the drug can drop the infection rate *β* to 0.1*β*, and the simulated results are shown in Figure .4. These results show that the earlier to give an effective drug to a serve patient, the better to relieve symptoms.

**Figure 4:**
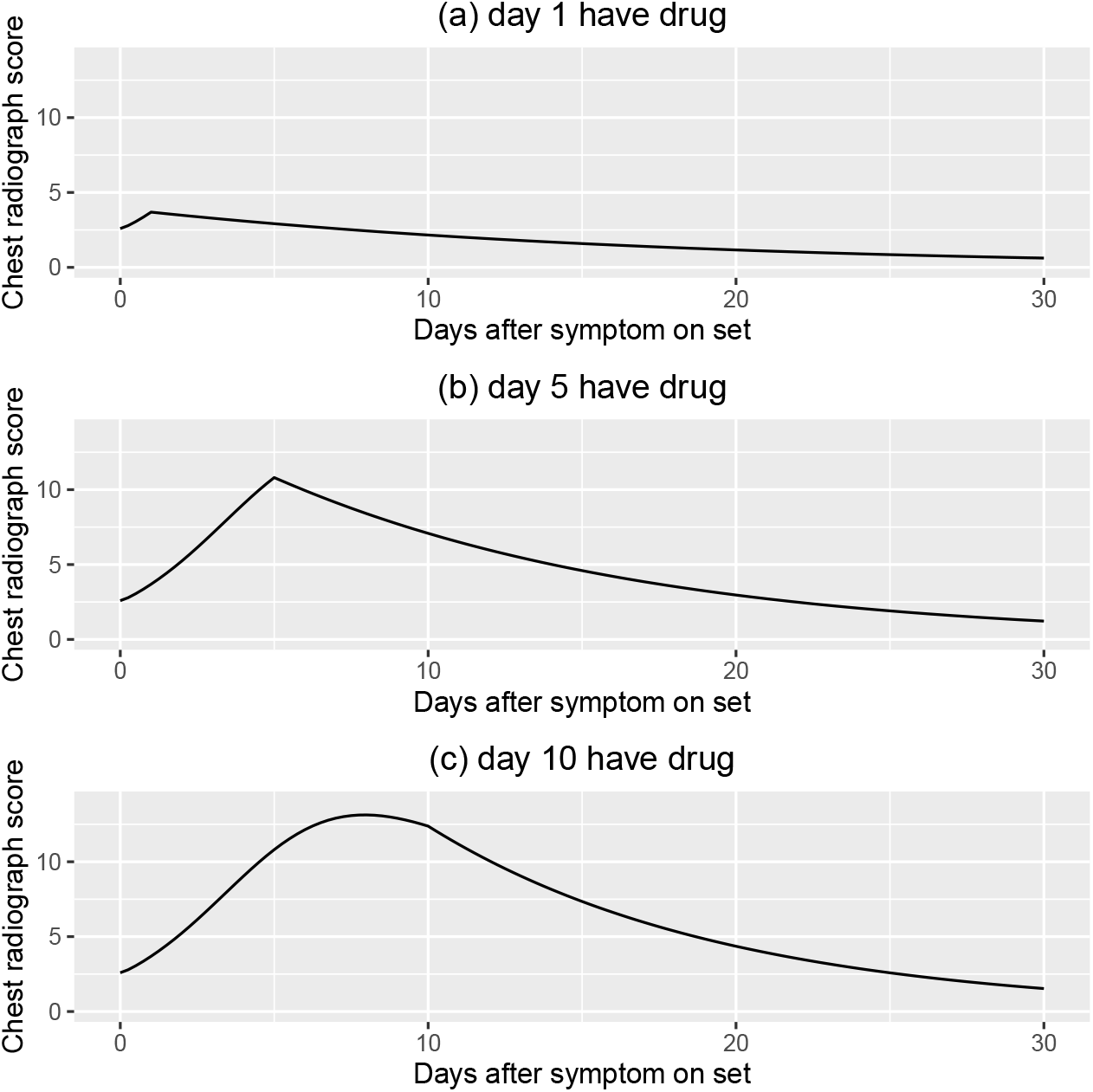
The simulated result of chest radiograph score of patients with COVID-19 and the beginning time of having drug is (a) the first day, (b) the fifth day, and (c) the tenth day after symptom onset. In this simulation, the drug effect is assumed to decrease the infection rate *β* to 0. 1 *β*.

## 4 Discussion

Based on the chest radiograph data, we estimate the parameters and basic reproductive number of the model (1). The *R*_0_ of SARS-CoV-2 in host growth is 3.79, which is higher than HCV (2.3, [2]), but lower than the flu virus (23, [10]). Comparing the estimated parameters of SARS-CoV-2 with MERS coronavirus, the MERS have a higher virulence to infect the pulmonary epithelial cells (larger *R*_0_) and with a small initial value of infected cell. These may esplain why the MERS have a higher mortality and smaller incubation period [6, 13]. By the PRCC method, the sensitivity analysis is also done on the *R*_0_, and the positive correlation of *E_p_*(0) with *R*_0_ may give a possible reason why the baby patients are less likely to die of this virus. We also analyze the drug therapy and its effect on the virus growth, and the result illustrates that early medication is effective for treatment.

The methods we used in this study are following the previous works by Miao et. al [10,11]. This is the first time for this methodology to be used in the COVID-19 and our work shows some new results of the within-host properties of this virus. All of the source codes are available at the GitHub (https://github.com/ChentongLi/SARS-CoV-2_viral_kinetic). Anyone could use these codes to estimate and forecast the chest radiograph score of the patients.

The major limitation of this study is that the chest radiograph score is not the real data of the infected pulmonary epithelial cells but just an approximation. Moreover, we assume that the immunne effect on viruses and infected epithelial cells are constant values. This is because the data of antibody and effective CD8 cells of that virus are rarely known. If we could get more accurate data, the results will be much better.

We believe this study could give some help to the diagnosis and treatment of the COVID-19. For other disease that infect the lung, this method we believe also could be used to do analysis.

## References

[1] Sebastian Bonhoeffer, Robert M May, George M Shaw, and Martin A Nowak. Virus dynamics and drug therapy. Proceedings of the National Academy of Sciences, 94(13):6971–6976, 1997.

[2] Swati DebRoy, Benjamin M Bolker, and Maia Martcheva. Bistability and long-term cure in a within-host model of hepatitis c. Journal of Biological Systems, 19(04):533–550, 2011.

[3] Or Dieckmann and JP Heesterbeek. Mathematical epidemiology of infectious diseases, 2000.

[4] Gunter Kampf, Daniel Todt, Stephanie Pfaender, and Eike Steinmann. Persistence of coronaviruses on inanimate surfaces and its inactivation with biocidal agents. Journal of Hospital Infection, 2020.

[5] Ha Youn Lee, David J Topham, Sung Yong Park, Joseph Hollenbaugh, John Treanor, Tim R Mosmann, Xia Jin, Brian M Ward, Hongyu Miao, Jeanne Holden-Wiltse, et al. Simulation and prediction of the adaptive immune response to influenza a virus infection. Journal of virology, 83(14):7151–7165, 2009.

[6] Qun Li, Xuhua Guan, Peng Wu, Xiaoye Wang, Lei Zhou, Yeqing Tong, Ruiqi Ren, Kathy SM Leung, Eric HY Lau, Jessica Y Wong, et al. Early transmission dynamics in wuhan, china, of novel coronavirus-infected pneumonia. New England Journal of Medicine, 2020.

[7] Tao Liu, Jianxiong Hu, Min Kang, Lifeng Lin, Haojie Zhong, Jianpeng Xiao, Guanhao He, Tie Song, Qiong Huang, Zuhua Rong, et al. Transmission dynamics of 2019 novel coronavirus (2019-ncov). 2020.

[8] Hongzhou Lu. Drug treatment options for the 2019-new coronavirus (2019-ncov). BioScience Trends, 2020.

[9] Simeone Marino, Ian B Hogue, Christian J Ray, and Denise E Kirschner. A methodology for performing global uncertainty and sensitivity analysis in systems biology. Journal of theoretical biology, 254(1):178–196, 2008.

[10] Hongyu Miao, Joseph A Hollenbaugh, Martin S Zand, Jeanne Holden-Wiltse, Tim R Mosmann, Alan S Perelson, Hulin Wu, and David J Topham. Quantifying the early immune response and adaptive immune response kinetics in mice infected with influenza a virus. Journal of virology, 84(13):6687–6698, 2010.

[11] Hongyu Miao, Xiaohua Xia, Alan S Perelson, and Hulin Wu. On identifiability of nonlinear ode models and applications in viral dynamics. SIAM review, 53(1):3–39, 2011.

[12] NCBI. The innate and adaptive immune systems. https://www.ncbi.nlm.nih.gov/books/NBK279396/#_i2255_theadaptiveimmunesys_.

[13] Myoung-don Oh, Wan Beom Park, Pyoeng Gyun Choe, Su-Jin Choi, Jong-Il Kim, Jeesoo Chae, Sung Sup Park, Eui-Chong Kim, Hong Sang Oh, Eun Jung Kim, et al. Viral load kinetics of mers coronavirus infection. New England Journal of Medicine, 375(13):1303–1305, 2016.

[14] Feng Pan, Tianhe Ye, Peng Sun, Shan Gui, Bo Liang, Lingli Li, Dandan Zheng, Jiazheng Wang, Richard L Hesketh, Lian Yang, et al. Time course of lung changes on chest ct during recovery from 2019 novel coronavirus (covid-19) pneumonia. Radiology, page 200370, 2020.

[15] The Novel Coronavirus Pneumonia Emergency Response Epidemiology Team. The epidemiological characteristics of an outbreak of 2019 novel coronavirus diseases (covid-19) in china. Chinese Journal of Epidemiology, 41(2):145–151, 2020.

[16] Wendi Wang and Xiao-Qiang Zhao. Threshold dynamics for compartmental epidemic models in periodic environments. Journal of Dynamics and Differential Equations, 20(3):699–717, 2008.

[17] WHO. Coronavirus disease (covid-19) outbreak. https://www.who.int/emergencies/diseases/novel-coronavirus-2019.

[18] Joseph T Wu, Kathy Leung, and Gabriel M Leung. Nowcasting and forecasting the potential domestic and international spread of the 2019-ncov outbreak originating in wuhan, china: a modelling study. The Lancet, 2020.

